# Droplets Bumpers as Mechanical Sensors for Cell Migration Under Confinement

**DOI:** 10.1101/037275

**Authors:** D. Molino, S. Quignard, C. Gruget, F. Pincet, Y. Chen, M. Piel, J. Fattaccioli

**Affiliations:** École Normale Supérieure - PSL Research University, Département de Chimie, 24 rue Lhomond, F-75005 Paris, France; Sorbonne Universités, UPMC Univ. Paris 06, PASTEUR, F-75005, Paris, France.; CNRS, UMR 8640 PASTEUR, F-75005, Paris, France; Laboratoire de Physique Statistique, Ecole Normale Supérieure, Université Pierre et Marie Curie, Université Paris Diderot, Centre National de la Recherche Scientifique UMR8550, 24 rue Lhomond, Paris 75005, France.; Institut Curie, CNRS UMR 144, 26 rue d’Ulm, 75005, Paris, France

## Abstract

Mechanical cell forces play determinant roles in cell growth and tissue development. Cell migration is a relevant context to explore single cell forces, since mechanotransduction and cell morphology are dynamically and biochemically co-regulated in order to modulate cell shape changes minute to minute. In this work we present the design of a hybrid microdevice made from a set of parallel PDMS microchannels in which oil emulsion droplets are introduced to be used as biocompatible and deformable cell forces transducers upon HL-60 human leukemia cell line migration. We show that the mechanical stress exerted by the cells can be measured from the analysis of the droplets deformations at the single cell level. Finally we show that acto-myosin contraction plays a role in the cell ability to cross obstacles in confinement conditions.

## Introduction

The ability of immune cells to migrate within narrow spaces is a critical feature involved in various physiological processes from immune response to metastasis. For instance, cells such as neutrophils are required to migrate within constrictions that are much smaller than their own diameter, such as small capillaries (*ca*. 2 μm)^1^. This ability greatly depends on the mechanical properties of the cells, since, for example, an increase of the cell stiffness has been shown to increase their retention in blood capillaries, eventually leading to inflammation^2, 3^.

The intracellular machine acting during cell migration is complex and involves cell surface adhesion molecules, cytoskeleton and its associated molecular motors, actin-plasma membrane interface, nucleoskeleton and mechano-transduction feedbacks^4–7^. *In vivo* assays of cell migration require the use of sophisticated microscopic techniques on live animals, such as *intravital microscopy*^8^, that are technically challenging. For the sake of simplicity and also to permit biophysical modeling of the migration processes, several *in-vitro* techniques have been developed^9^ as the modified Boyden chamber^10^ or transwell assay^11^ that provide end-point data but no information on cell behavior, especially its mechanics, between the start and conclusion of the experiment. However, thanks to the engineering of techniques based on the analysis of deformable substrates such as thin silicon membranes^12^, 2D and 3D gels^13–15^ or flexible pillars^16, 17^, our understanding about the stress generation pathways involved in cell migration has largely improved.

In confinement conditions, studies performed with PDMS microdevices have shown that nuclear deformability is one of the limiting factors that slows down and even impedes the ability of cells to migrate within microfabricated constrictions^18–21^. The mechanical rigidity of the fabrication materials such as PDMS^22^ limits the collection of quantitative data related to the physical stress a cell is able to produce when crossing a constriction during a migration event, thus pushing for the development of microdevices having softer actuation elements with mechanical properties comparable to those of cells^23^.

As an alternative to polymers or hydrogels that are more commonly used when soft substrates are needed^24^, we propose in this study to use oil-in-water emulsion droplets as *in vitro* mechanical sensors during cell migration, since their stiffness has been shown to be comparable to the one measured for cells^25^. Hence we developed a hybrid microchip made of parallel PDMS channels in which oil droplets, with sizes comparable to cells, are sparsely distributed and serve as deformable obstacles that migrating cells have to squeeze to explore their environment. Since the shape of a droplet is set by the interplay between the interfacial tension and the mechanical stress field acting on it^26, 27^, a simple microscopic analysis of the deformation of the droplet shape over time brings quantitative information on the mechanical stress that cells are exerting on it.

After a description of the fabrication of the microdevice, we show that HL-60 cells are able to cross and squeeze the obstacles while deforming their nucleus. We then describe the quantitative analysis procedure of the droplet deformation and we quantify the mechanical stress exerted by a cell on a droplet during crossing events. We finally show that the ability of a cell to go through droplet obstacles is actomyosin dependent. Our system hence provides a simple *in vitro* tool to explore by live imaging the mechanic and the molecular machinery necessary for a cell to infiltrate tissues.

## Materials and Methods

**Emulsion droplets fabrication and staining.** Oil droplets are made from soybean oil (Sigma-Aldrich, St. Louis, MO, USA). Briefly, soybean oil was dispersed and emulsified by hand in an aqueous continuous phase containing 15 % w/w of Poloxamer 188 block polymer surfactant (CRODA, East Yorkshire, UK) and 1 % w/w sodium alginate (Sigma-Aldrich, St. Louis, MO, USA) at a final oil fraction equal to 75%. The rough emulsion was sheared in a Couette cell apparatus at a controlled shear rate of 110 rpm as described by Mason *et al*. ^28^. For storage and handling purposes the emulsion are diluted to an oil fraction of 60 % w/w with 1 % w/w of poloxamer 188 in the continuous phase and stored at 12°C in a Peltier-cooled cabinet.

To stain droplets with Nile Red (Sigma-Aldrich, St. Louis, MO, USA), a red lipophilic dye, the droplets suspension is washed and resuspended in cell growth complete media containing 10μM of Nile Red. Size distribution of the emulsion droplets was measured by brightfield microscopy and image analysis.

**Cell culture handling**. HL-60 expressing GFP-Actin (kindly provided by Guillaume Charras, from UCL, UK) were grown in RPMI media supplemented with 15% fetal bovine serum, 50 mM Hepes, 2mM L-Glutamine, 10 units/penicillin and 10 mg/ml streptomycin (Life Technologies, California, USA), in a 5% CO2-humidified atmosphere at 37°C. Cells were passaged to 0.15 million cells per mL when a maximal density of 1-2 million cells per mL was reached (*ca*. every 2-3 days). Passages were done in a total volume of 10 mL pre-warmed culture medium, in 25 cm^2^ cell culture flasks with 0.2 μm filter cap (Nunc™, Roskilde, Denmark). HL60 cells were differentiated with 1.3 % v/v DMSO for 6 days without antibiotics ^29^. For all migration experiments cells were loaded in microchip in self-conditioned media.

**Cell staining and treatments**. For nuclei staining cells where treated with 0.5 μg/ml Hoechst 33342 (Life Technologies, Carlsbad, California, USA) for 30 minutes at 37°C. For Y-27632 treatment, cells were loaded in chip and incubated for 4h with 10 μM Y-27632, in self-conditioned media. For quantification of the Y-27632 effect only channels containing less than 4 drops where taken into account. In each channel the number of cells localized before the first droplet or after where counted along total distance of 700 μm from the loading well.

**PDMS microchips with cells and drops**. Channel based PDMS chips where fresh replicates from epoxy mold and glowed on Fluorodish F35 (from WPI, Sarasota, FL 34240, USA) as described by Vargas *et al*.^30^. After 30 min under vacuum soybean drops pre-incubated in cell growth media were charged into loading wells at a 1:300 dilution. Cells were loaded immediately after at concentration of 10^5^ cells per well of 2,5 mm diameter. For all migration experiments cells were loaded in the microchip within their self-conditioned media.

**Microscopy**. Live cells were imaged on a LSM 710 (Zeiss) confocal microscope, using a 405 nm laser diode exciting Hoechst 33342, a 488 nm argon laser line exciting GFP and a 561 nm diode laser line exciting Nile Red. Emission was detected between 410 and 480 nm for Hoechst 33342, 495–530 nm for GFP and 565–max nm for Nile Red. Live imaging studies were made at 37°C in differentiating media. Acquisition was made in channel-separated mode and with a line-scanning mode, with a line average of 2 and an 8-bit dynamic range. Images were analyzed using the softwares Fiji/Image J ^31^ and Mathworks Matlab softwares.

**Interfacial tension measurement**. To measure the interfacial tension of soybean oil droplets we used the micropipette-aspiration method that has been described in detail in a previous article^32^. Micropipettes were made from 1 mm borosilicate glass-tube capillaries (Harvard Apparatus, USA) that were pulled in a pipette puller (P-2000, Sutter instrument Co., USA) to tip diameters in the range of 2 to 3 μm. An oil hydraulic micromanipulator (Narishige, Japan) allowed for pipette positioning and micromanipulation. The pipette was connected to water reservoirs that could be translated vertically to apply precise suction pressures.

**Statistical analysis**. Data were processed first with Wilcoxon test to evaluate data distributions. Data with Gaussian distributions were validated with paired t-test, non-Gaussian distributed sets of data were evaluated with Mann-Whitney. Graphs and statistics were obtained using GraphPad Prism software

## Results

**Description of the hybrid microdevice**. We developed a hybrid microfluidic device (**Figure 1A**) inspired by the micro-channel based assay that has been used so far to study dendritic cells migration under confinement^21^. The PDMS device consists of two circular loading chambers (2.5 mm diameter) connected by 25 parallel rectangular channels (width: 14 μm, height: 8 μm, length: 2 mm). In each channel, soybean oil emulsion droplets with dimensions comparable with those of the channels are randomly distributed and serve as deformable obstacles on the path of migrating immune cells.

**Figure 1:**
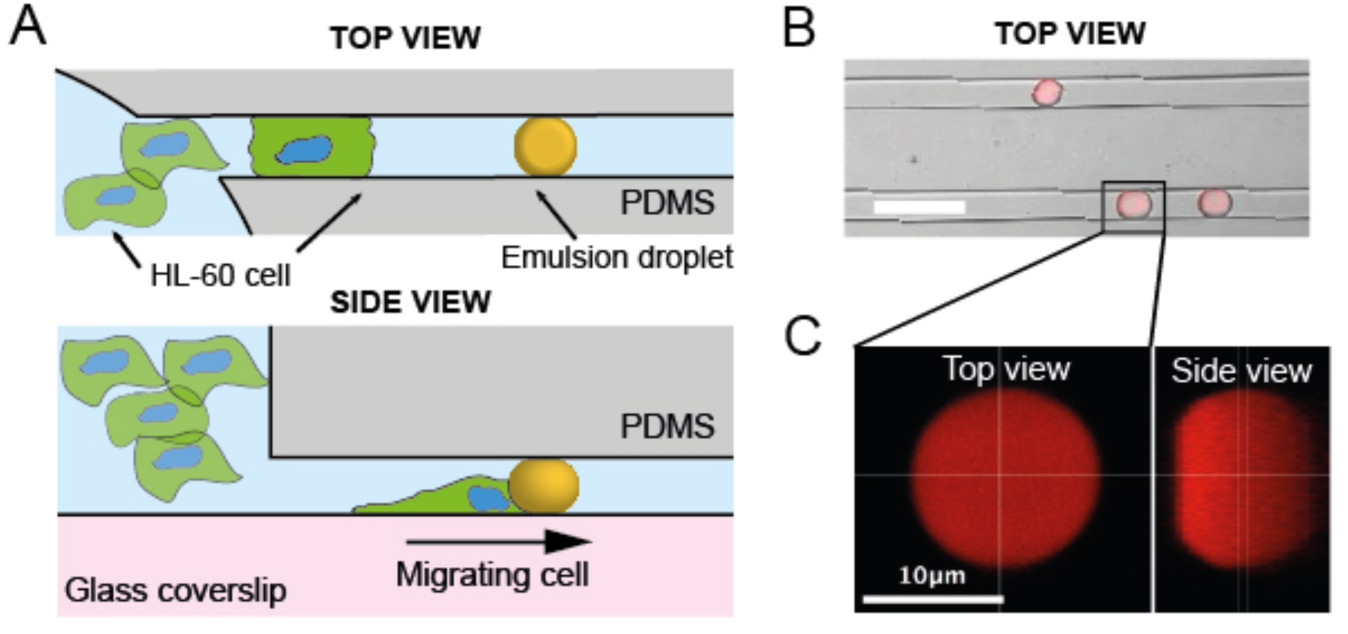
(A) Schematic view of the hybrid microdevice made from two circular loading chambers (diameter: 2.5 mm) connected by parallel rectangular channels (width: 14 μm, height: 8 μm). Soybean oil droplets are inserted within the channels and make a deformable obstacle to the migration of HL-60 cells. (B) Confocal microscopy picture of PDMS channels containing oil droplets stained with Nile Red dye. Scale bar: 30 μm. (C) Top view and 3D reconstructed side view of a droplet blocked in a micro channel. Scale bar: 10 μm

Emulsions are colloidal liquid-liquid metastable suspensions stabilized by a surfactant monolayer^33^ that have already shown theie biocompatibility in a biophysical context^34, 35^. To fabricate the droplets, we manually shear soybean oil in an aqueous solution of a polymeric surfactant (poloxamer 188) to stabilize the emulsion and a viscosifier (sodium alginate) to increase the viscoelasticity of the continuous phase and ease the fragmentation. The crude, polydisperse emulsion is then sheared in a Couette cell apparatus, following the method developed by Mason *et al*.^28^, to obtain a quasi-monodisperse, 13 ± 2 μm diameter emulsion (**Figure S1A**). Prior to the migration experiments, the continuous phase is replaced by normal cell culture medium thanks to several centrifugation/rinsing steps.

After fabrication and mounting of the PDMS on a small Petri dish with a glass bottom, a dilute suspension of deformable soybean oil droplets is injected into one of the loading well at a concentration allowing the droplets to be sparsely distributed in the channels (**Figure 1B**). The size of the droplets was chosen to be slightly larger than the smallest dimension of the channels, so drops with a diameter bigger that 8 μm are squeezed horizontally and hence remain immobilized into the device. 3D reconstructions of Nile Red-stained droplets into the channels (**Figure 1C**) show that droplets with a diameter larger than the channel width are pancake-shaped, the top and bottom interfaces being flat thanks to pressure exerted by the PDMS walls in the *z* dimension (**Figure S2**). We confirmed the shape of the droplets in this size range by numerical simulations (**Figure S1B**) made with the software Surface Evolver ^36^.

**Interfacial tension of the droplets**. To carefully evaluate the interfacial tension of the droplets suspended in the culture medium used for migration experiments, we used the micropipette aspiration method^32^. Upon aspiration by a very thin pipette, the droplet deforms and a spherical cap of radius R_c_ forms at the tip of the pipette (**Figure 2A**). At equilibrium, the value of R_C_ depends on the interfacial tension γ of the droplets, the radius of the droplet R_D_, the aspiration pressure ΔP and can be expressed as:

**Figure 2:**
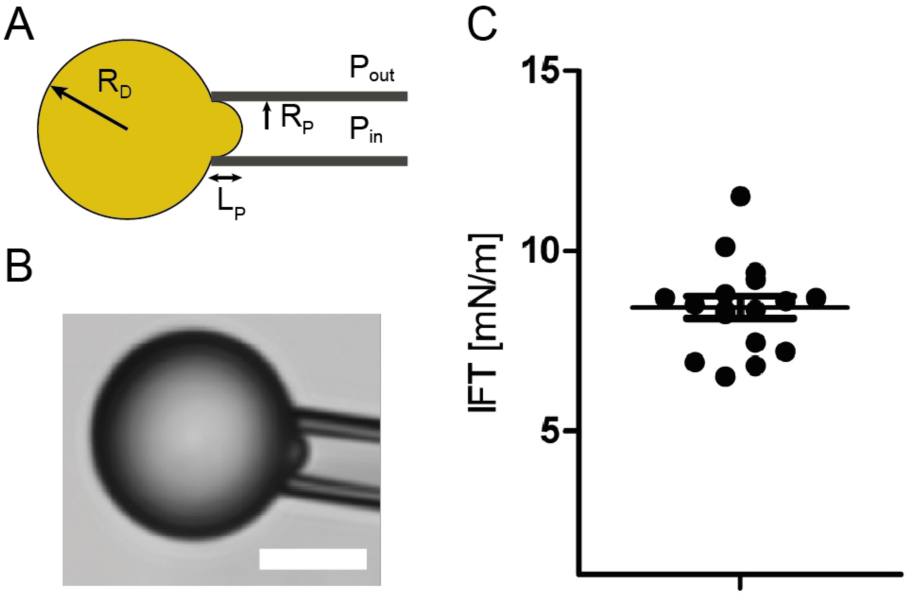
(A) Schematic view of the micropipette experiment used to measure the interfacial tension
of the droplets. R_P_ and R_D_ are respectively the pipette and the droplet radius. (B) Bright field microscopy image of the droplets aspirated by the micropipette. (C) Plot showing the interfacial tension (IFT) of soybean oil droplets, dots are independent droplets (N=17). Mean interfacial tension is 8.3 ± 1.26 mN.m-1.

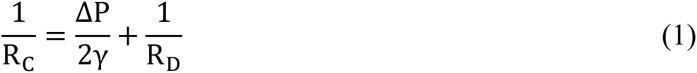

The aspiration ΔP corresponds to the pressure difference between the inside of the pipette and the external pressure. γ and R_D_ are constant throughout the experiment. Hence, varying ΔP induces changes in R_C_: the larger ΔP, the smaller R_C_. During the course of a measurement, ΔP is slowly increased and R_C_ decreases until it reaches the radius of the pipette R_P_. Up to that critical aspiration ΔP_c_, little change is observed in the geometry of the system. As soon as ΔP is larger than ΔP_c_, R_C_ becomes smaller than R_P_, which results in the sudden entry of the oil droplet in the pipette. This provides a direct measurement of the surface tension of the droplet:

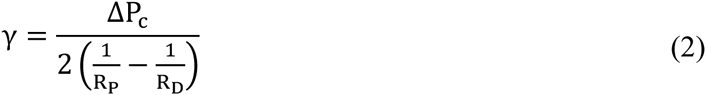

Each droplet can be blown out and aspirated again several times, allowing averaging the measurement and refining the interfacial tension value. The interfacial tension of the soybean oil droplets suspended into the differentiating media, at 37°C, is equal to 8.4 ± 1.2 mN.m^-1^ (**Figure 2B**).

**HL-60 migration in the microdevice**. Cells of the immune system, such as HL-60, represent a simple model system to study cell migration without the need to derive cells from primary tissue ^37^. The capability of differentiated HL-60 to bypass endothelial barriers renders this cell type particularly suitable for studying interstitial cell migration *in vitro* ^38–40^.

After insertion of the droplets in the microchannels, neutrophil-like cells derived from DMSO differentiated^29^ HL-60 cells expressing GFP-actin are seeded in one of the loading well and the microchip is put at rest in a culture incubator to allow the cells to settle and recover their motility. After around 2 hours, cells spontaneously start entering channels where droplets are inserted, as observed by live imaging (**Movie S1**). HL-60 cells migrating within a microchannel move in the forward and backward direction relative to the loading well.

When a cell encounters a droplet within the microchannel, its behaviour towards the obstacle strongly depends on the size of the droplet. For droplets whose diameter is smaller than the width of the channel, HL-60 cells migrate while displacing the droplets with them over long distances, as shown in **Figure 3A** and **Movie S2**. In the case of droplets that are as large as and larger than the microchannel width, cells are not able anymore to move the droplets while migrating but rather cross the deformable obstacle (**Figure 3B**). Although cells could potentially go through the droplets along any of the four walls of the microchannel, they actually never pass on the top and bottom flat sides of the pancake-shaped droplet but move along the rounded parts instead, which definitely ease the timelapse observation (**Movie S1).**

**Figure 3:**
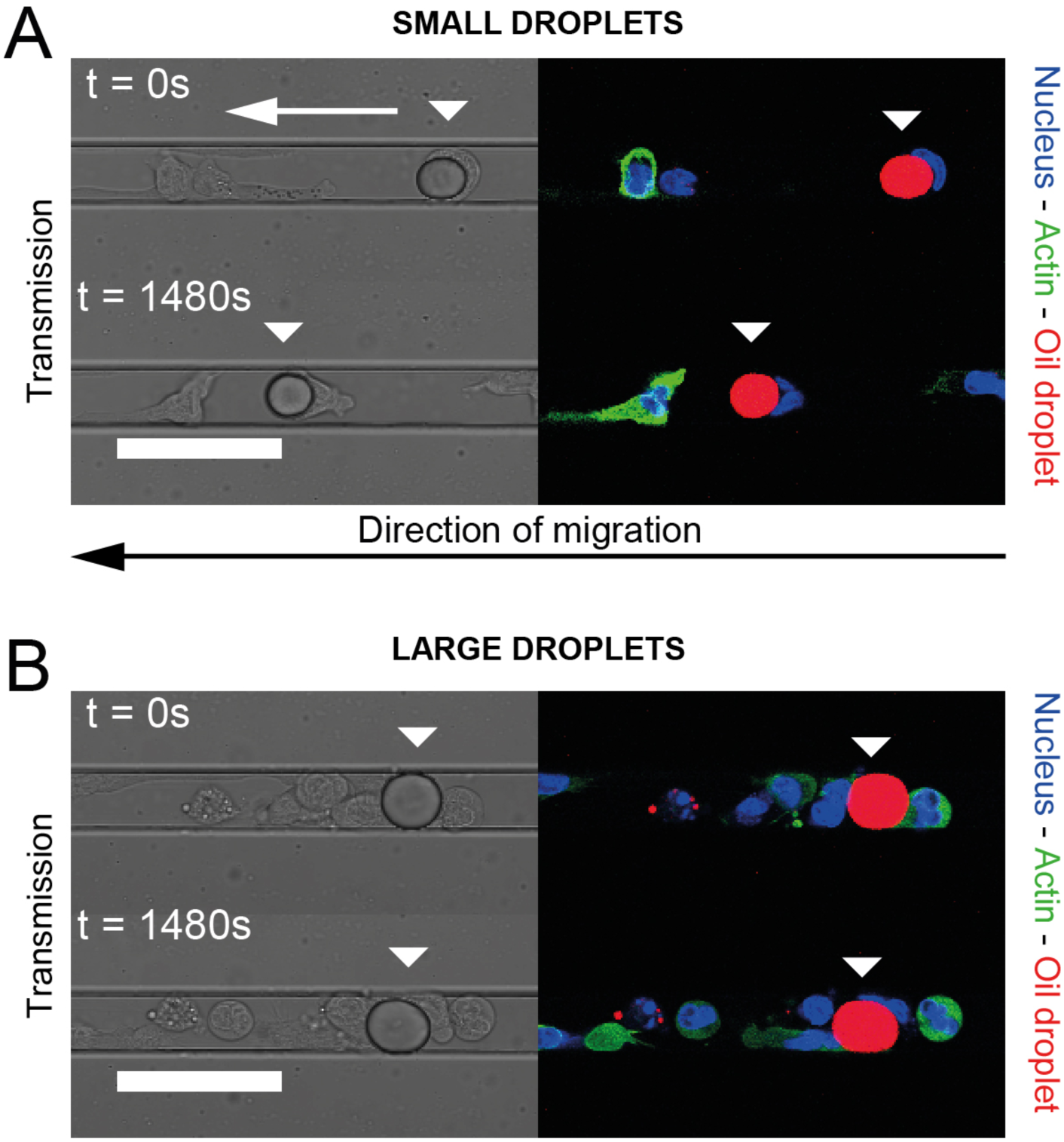
Time-lapse recording of the behavior of HL-60 cells (Hoechst 33342: blue, Actin-GFP: green) migrating along a channel and encountering a small (A) and a large (B) soybean oil droplet (Nile Red: red), the latter being large enough to be in contact with the walls of the PDMS microchannel. The relative positions of the arrows indicate the displacement of the droplets due to the cells. Scale bar: 40 μm.

**Figure 4A** shows a time-lapse recording of a crossing event recorded in the focal plane located in the middle of the microchannel. We see that both cell and droplet are deformed during the encounter as a result of the mechanical stress applied by the cell on the droplet to go through it. While the nucleus of the neutrophil has a rounded aspect before and after the encounter with the droplet, the nucleus gets squeezed and elongates during the crossing. Contrary to the front part of the cell actin cortex that can cross the obstacle without deforming it, the maximal deformation of the droplet is observed when the cell nucleus is going from one side to the other of the droplet (**Figure 4B**). The droplet, circular in its resting state, becomes pear-shaped when the cell is pushing on it, and finally recovers its resting shape when the cell moves away from it (**Figure 4A**).

**Figure 4:**
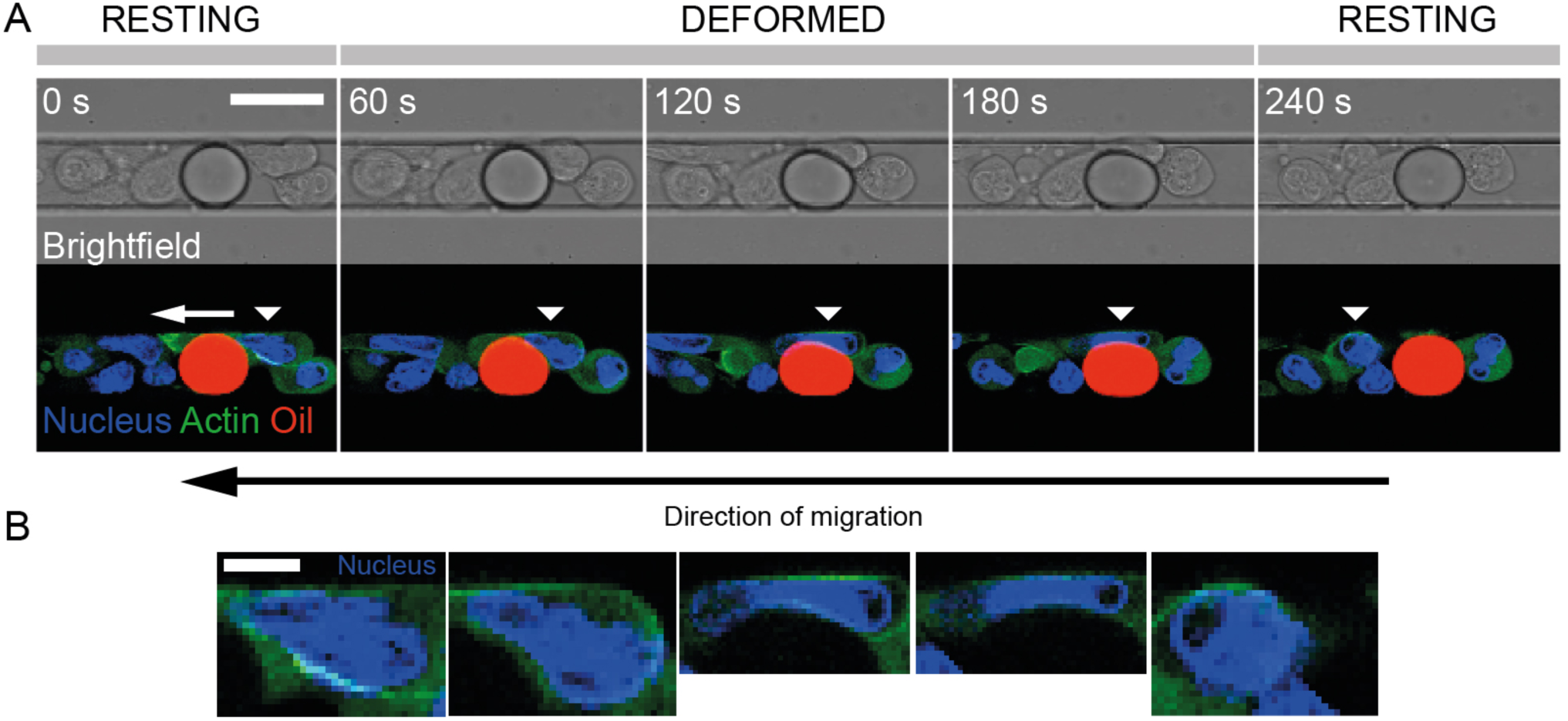
(A) Time-lapse recording of an HL-60 cell (Hoechst 33342: blue, Actin-GFP: green) migrating along a channel and squeezing a soybean oil droplet (Nile Red: red). The moving cell is identified with a white triangular mark. Scale bar: 15 μm. (B) Corresponding enlarged pictures of the nucleus of the migrating cell in (A). Scale bar: 5 μm.

**Mechanical description of the droplet shape**. The shape of an oil droplet at the equilibrium is governed locally by the interplay between its interfacial tension γ and the mechanical stress, homogeneous to a pressure, exerted by the environment on the droplet.

As a consequence of the existence of an interfacial tension γ, according to the Young-Laplace equation ^41^, the local shape of the interface of a droplet confined in a microchannel with an initial radius *R*_0_ obeys the general relationship, in spherical coordinates:

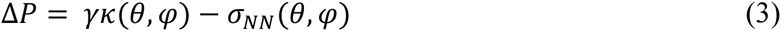

where Δ*P* = *P_in_* – *P_out_* is the pressure excess across the emulsion interface, *κ (θ, φ)* the mean curvature of the interface and *σ_NN_ (θ, φ)* the normal mechanical stress acting on the droplet.

The mean curvature *κ (θ, φ)* writes as

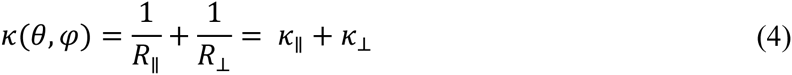

where *R*_∥_ and *R*_⊥_ are the two principal radii of curvature.

In absence of hydrodynamic flow, *P_out_* from **Equation 4** is a constant equal to the hydrostatic pressure within the microchannel. Moreover, **Figure 4** and **Figure S3** show that the HL-60 cells do not surround completely the droplet, meaning that a part of the oil interface is always in contact with the aqueous culture medium. As the inner pressure *P_in_* is homogeneous within the droplet, its value is hence set by the oil/medium rather by the oil/cell membrane interfacial tension, which makes the excess pressure *ΔP* of the droplet a constant at each time point of the experiment.

The local mechanical stress *Δσ_NN_* exerted by a cell can thus be measured according to the variation of the principal curvatures between the deformed and the resting state of the droplet within the contact area with the cell:

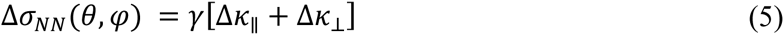

In its resting state, the principal radii of curvature are respectively measured in-plane (*R*_∥_) and out-ofplane (*R*_⊥_) to the focal plane of observation corresponding to the **Figure 4**. In the following, we assume that the focal plane still remains a plane of principal curvature also in the case where the oil droplet gets deformed during the encounter.

Despite the large in-plane deformation of the droplet upon migration of a cell shown in **Figure 4**, the out of plane deformation is almost inexistent (**Figure S4**), meaning that the corresponding curvature *κ*_⊥_ doesn’t change over time and its variation can be neglected in the mechanical stress computation. A correct estimation of the stress *σ_NN_* can hence be derived from the value of the interfacial tension and the in-plane curvature variation between resting and deformed droplet state:

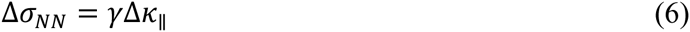

**Segmentation and numerical computation of the in-plane curvature κ_∥_**. To measure the curvature *κ*_∥_ within the focal plane of observation, we proceed to a segmentation of the oil droplets and to their conversion in a shape that can be numerically processed to extract geometric data of interest (**Figure 5**). Briefly, z stack images of the droplets stained with Nile Red were slightly denoised with a median filter to homogenize their aspect and ease the segmentation. Then, we used an active contour (*snakes*) routine to record the (x,y) Cartesian coordinates of the droplet interface. For the active contours to take a full account of the intensity gradient on the border of the droplets and ultimately increase the accuracy of the segmentation, the pictures were first scaled 10 times without interpolation to artificially subdivide each pixel of the original image in a 10x10 pixel grid (**Figure 5A**). The pixelation of the contour coordinates was then suppressed by the application of a low-pass 1D FFT filtering both on the x and y coordinates, to get the interpolated model droplet. We finally computed the local analytical curvature of the droplets shape from the (x,y) Cartesian filtered data, as shown on (**Figure 5B**).

**Figure 5:**
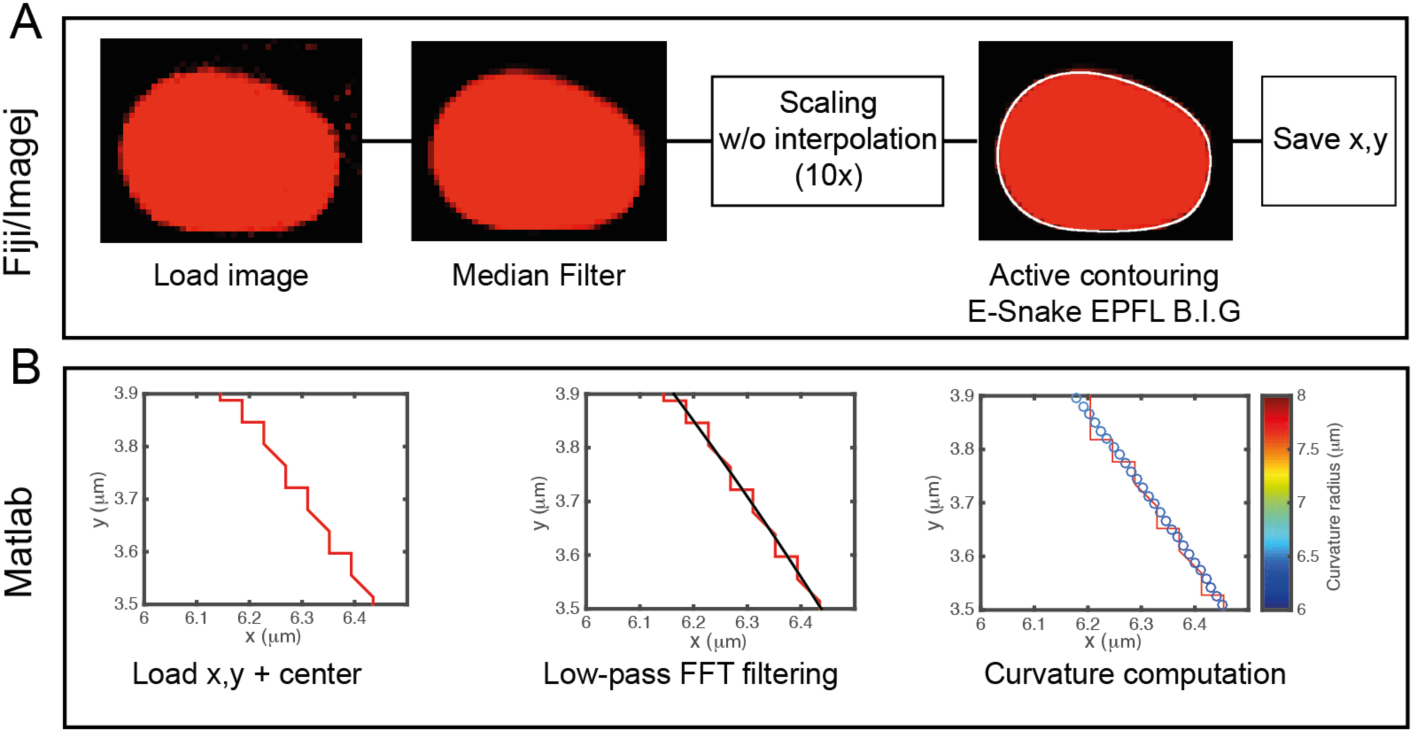
Workflow of the droplet segmentation routines. (A) Fiji/ImageJ segmentation procedure: confocal microscopy pictures of the droplets are first slightly denoised with a median filter then pictures are scaled 10 times and we used an active contour (*snakes*) routine to record the (x,y) Cartesian coordinates of the droplet interface after a visual comparison with the experimental picture. (B) Matlab curvature computation: the pixelation of the contour coordinates is suppressed by the application of a low-pass FFT filter. The local analytical curvature of the droplets shape is computed from the filtered data. Pictures show a zoomed area of the droplet contour.

**Mechanical stress exterted by the cells**. From time-lapse recordings similar to the one shown in **Figure 4**, we selected a frame where the droplet is resting, either before or after the crossing by a cell, and a frame where the droplets is squeezed and pear-shaped as a cell is pushing on it. Using the segmentation and computation routine described above, we obtain the shape outlines shown on **Figure 6A,B** for resting and deformed oil droplet. For the sake of simplicity, the shape outlines are color-encoded with respect to the local radius of curvature. As expected when considering the simulations in **Figure S1B**, **Figure 6A** shows that the droplet in the resting state is globally rounded and slightly flattened at locations where it touches the walls of the microchannel. When the cell pushes on the droplet to go though it, the radius of curvature increases in the region of contact with the cell and on the bottom part of the droplet. **Figure 6C** shows the evolution of the local analytical curvature along angular coordinates that correspond to the upper part of the droplet. The maximal mechanical stress is calculated at the location where the curvature of the constrained droplet is minimal (**Figure 6A,B** - arrow). As shown on **Figure 6D**, measurements performed on a sample of independent droplets (N= 8) give a numerical value Δ*σ_NN_* equal to 500 ± 100 Pa, which corresponds to a mechanical stress Δ*σ_NN_* = 500 ± 100 pN. μm^-2^.

**Figure 6:**
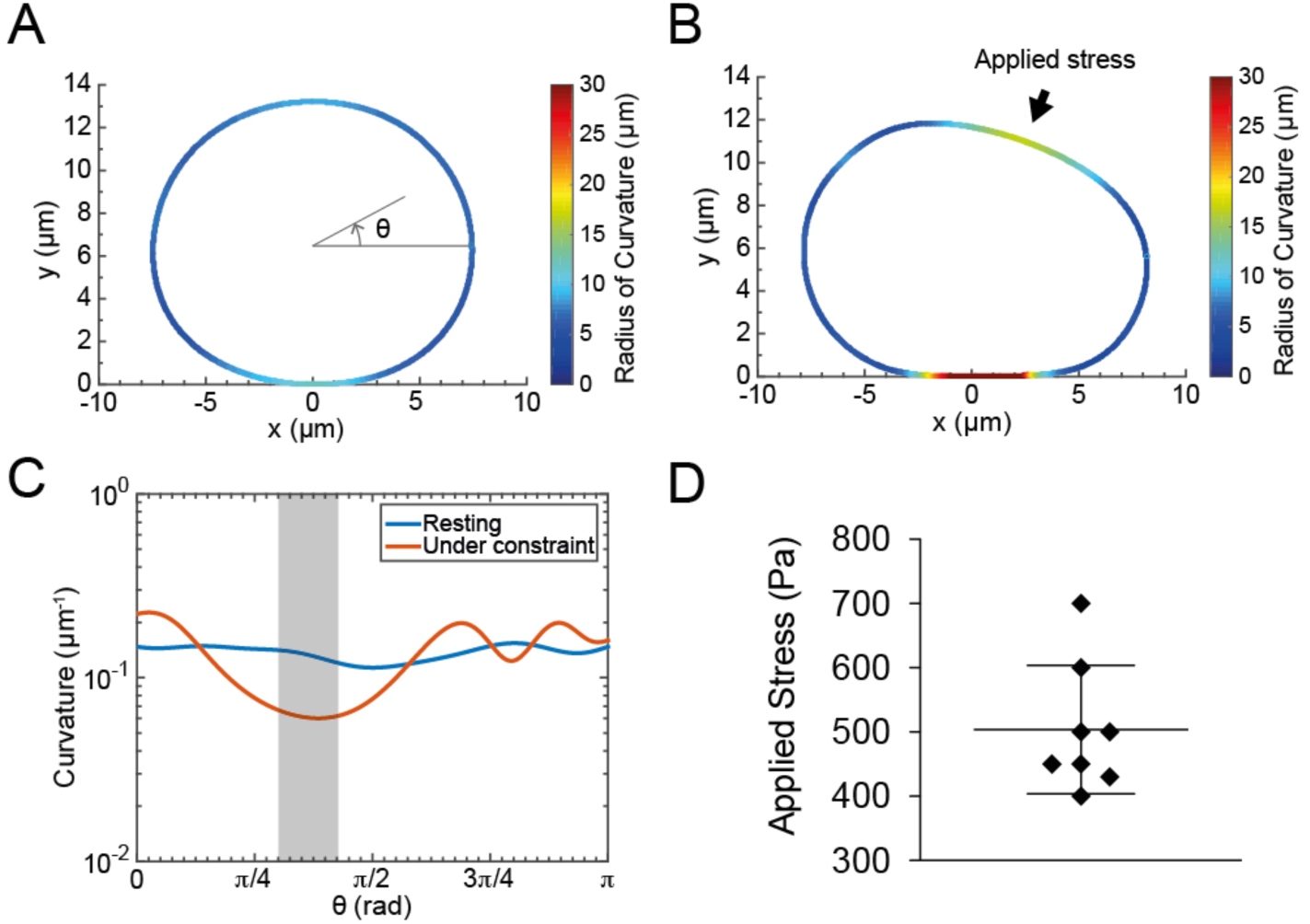
Resting (A) and constrained (B) droplet shape after segmentation and computation of the local radius of curvature. The shape outlines are color-encoded with respect to the local radius of curvature. (C) Analytical curvature as a function of the angular position along the droplet profile in the resting (blue) and constrained state (red). Only the values corresponding to the upper half part of the droplet are plotted. The grey area corresponds to the area where the mechanical stress applied by the cell is maximal. (D) Measured values of the mechanical stress exerted by the cells (N=8).

**HL-60 ability to go through a droplet is an actomyosin dependent process**. To assess the involvement of actomyosin contractility in HL-60 migration in the presence of droplets, we used the Y-27632 drug, which block ROCK1 phosphatase and in consequence myo-2 phosphorylation, necessary for actomyosin-dependent contraction ^42, 43^. After loading cells in one of the two inlets of the microchip, we counted for any microchannel filled with sparsely distributed droplets the number of cells localized either before or after the first drop in a range of 700 μm distance from the loading well (**Figure 7A**). 4 hours after cell loading we find that cells in control conditions are in average distributed equally before and after the droplets (**Figure 7B**), confirming the ability of the population of migrating HL-60 cells to go through the drop obstacles in an unbiased manner. Under Y-27632 condition however, most of the cells remain localized in the region between the loading well and the first droplet of the micro channel. The drug treatment thus interferes with the capability of the cell to bypass a drop.

**Figure 7:**
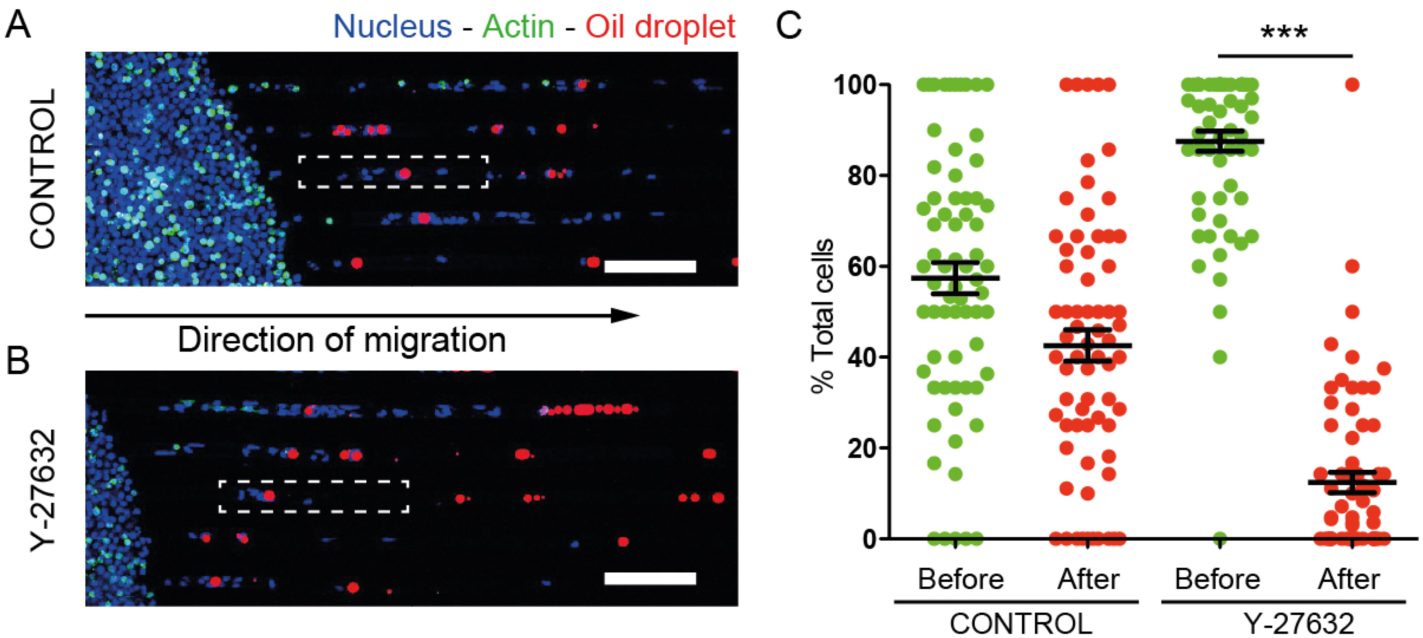
Comparison of the HL-60 spatial distribution along microchannels containing droplets in control and Y-27632 conditions. (A) Confocal microscopy mosaic images. Oil droplets are stained with Nile Red (red), cell nuclei with Hoechst 33342 (blue). Scale bar: 100 μm. (B). Percentage of cells localized before and after a droplet in a microchannel. N=68 channels for a total of 3 independent experiments. For control P > 0.01 (t-test), for Y-27632 treatment, P < 0.0001 (Mann-Whitney).

## Discussion

The microfluidic device described here allows monitoring cell migration and quantifying the ability of immune cells to cross obstacles of comparable size in real time, at single cell level. The chip design involves the trapping of quasi-monodisperse soybean oil-in-water droplets in PDMS microchannels within which cells migrate. When the droplets are large enough to be constricted by the four walls of the microchannel, they constitute immobile obstacles that migrating HL-60 cells have to squeeze to pass through. During this process, the cell nucleus gets strongly deformed and finally recovers its rounded shape when the cell reaches the other side of the obstacle. Besides the HL-60 cell line, we also used primary murine dendritic cells (**Figure S5**), which successfully deformed droplets during their crossing, showing that both cell types, albeit different, can exert a mechanical stress on the obstacles. From the physiological point of view, our experimental method is more related to the characterization of the invasiveness of a particular cell line in a crowded and confined environment than its migratory behaviour *stricto sensu*.

The choice of soybean oil as the formulation base was driven by the fact that it gives stable and biocompatible emulsions currently approved for pharmaceutical products^44^, but also because it has one of the lowest interfacial tension as compared to common formulation constituents as silicone or mineral oils^45^. This insures that a cell can squeeze a droplet sufficiently during the crossing to allow measuring a change in the droplet shape. From the analysis of their deformation and the measurement of their interfacial tension, droplets are hence used as mechanical sensors during migration events, and the mechanical stress exerted by the cell on the droplet is computed from the local curvature change at the locations where the cell contacts the droplet.

However, in presence of Y-27632, a drug able to inhibit the actomyosin contractility, our results show that differentiated HL-60 remain motile but are not able anymore to bypass droplets (**movie S3**), staying stuck on one side of the obstacles instead. These data suggest that the drug impairs cell forces generation related to the obstacle crossing.

The mechanical stress we measure from the analysis of the oil droplet deformation (Δ*σ_NN_* = 500 ± 100 pN.μm^-2^) is in the middle range of what has been measured so far for traction stresses of migrating cells, both in 2D and 3D conditions^13, 46-49^. Several studies clearly demonstrated that the organelle that limits the most cell deformation during migration in constrictions is the nucleus^2, 50, 51^, since it is the biggest and stiffest organelle of the cell^50^. From **Figure 4B**, we can estimate the smallest dimension the nucleus can reach when fully deformed is around 2-3 microns, which is in accordance with data from the literature. The mechanical stress value we provide here is thus related to the ability of a cell to deform an inert object of comparable stiffness in order to cross it, and can be seen as the amount of force the cell has to provide to create a space large enough so that its nucleus can flow from one side of the droplet to the other. Wether or not such stress depends on the rigidity of the obstacle is a question that should be addressed in the future.

## Conclusion

The microfluidic platform described here allows monitoring cell migration under confinement and measuring mechanical stresses exerted by immune cells during obstacle crossing in real time, over extended periods of time. This system is easy to handle and allows access to quantitative and qualitative information at the single-cell level on the involvement of intracellular players that are necessary to generate forces. We believe that this new system provides a simple tool to explore, by live imaging, the invasive ability of migrating cells in crowded environments.

## Acknowlegments

We thank Guillaume Charras (LCN) for having provided the Actin-GFP HL-60 cell lines; Ana-Maria Lenon-Dumesnil (Curie Institute) for having provided primary murine dendritic cells, J. Bibette (ESPCI) for lending us the Couette emulsifier; Nicolas Bremond (ESPCI) and Patrick Perrin (ESPCI) for their help regarding interfacial tension measurements; Thomas Boudier (UPMC, NSU) for his help on image segmentation.

This work has received support of "Institut Pierre-Gilles de Gennes » (Laboratoire d’excellence: ANR-10-LABX-31, “Investissements d’avenir”: ANR-10-IDEX-0001-02 PSL and Equipement d’excellence: ANR-10-EQPX-34).

## References

1 M. P. Wiedeman, Circ. Res., 1963, 12, 375–378.

2 G. P. Downey, D. E. Doherty, B. Schwab, E. L. Elson, P. M. Henson and G. S. Worthen, J. Appl. Physiol., 1990, 69, 1767–78.

3 G. S. Worthen, B. I. Schwab, E. L. Elson and G. P. Downey, Science, 1989, 245, 183–186.

4 V. Vasioukhin, C. Bauer, M. Yin and E. Fuchs, Cell, 2000, 100, 209–219.

5 F. Wang, M. Kovács, A. Hu, J. Limouze, E. V. Harvey and J. R. Sellers, J. Biol. Chem., 2003, 278, 27439–27448.

6 T. Lecuit, P.-F. Lenne and E. Munro, Annu. Rev. Cell Dev. Biol., 2011, 27, 157–184.

7 W. Engl, B. Arasi, L. L. Yap, J. P. Thiery and V. Viasnoff, Nat. Cell Biol., 2014, 16, 587–94.

8 D. Palm, K. Lang, B. Brandt, K. S. Zaenker and F. Entschladen, Semin. Cancer Biol., 2005, 15, 396–404.

9 C.-C. Liang, A. Y. Park and J.-L. Guan, Nat. Protoc., 2007, 2, 329–33.

10 S. Boyden, J. Exp. Med., 1962, 453–466.

11 D. R. Senger, C. a Perruzzi, M. Streit, V. E. Koteliansky, A. R. de Fougerolles and M. Detmar, Am. J. Pathol., 2002, 160, 195–204.

12 A. Harris, P. Wild and D. Stopak, Science, 1980, 208, 177–179.

13 W. R. Legant, J. S. Miller, B. L. Blakely, D. M. Cohen, G. M. Genin and C. S. Chen, Nat. Methods, 2010, 7, 969–71.

14 K. a. Beningo and Y. L. Wang, Trends Cell Biol., 2002, 12, 79–84.

15 X. Tang, A. Tofangchi, S.V Anand and T. a Saif, PLoS Comput. Biol., 2014, 10, e1003631.

16 J. L. Tan, J. Tien, D. M. Pirone, D. S. Gray, K. Bhadriraju and C. S. Chen, Proc. Natl. Acad. Sci. U. S. A., 2003, 100, 1484–1489.

17 O. du Roure, A. Saez, A. Buguin, R. H. Austin, P. Chavrier, P. Silberzan and B. Ladoux, Proc. Natl. Acad. Sci., 2005, 102, 2390–2395.

18 P. M. Davidson, J. Sliz, P. Isermann, C. Denais and J. Lammerding, Integr. Biol., 2015.

19 J. R. Lange, J. Steinwachs, T. Kolb, L. A. Lautscham, I. Harder, G. Whyte and B. Fabry, Biophys. J., 2015, 109, 26–34.

20 A. C. Rowat, D. E. Jaalouk, M. Zwerger, W. L. Ung, I. A. Eydelnant, D. E. Olins, A. L. Olins, H. Herrmann, D. A. Weitz and J. Lammerding, J. Biol. Chem., 2013, 288, 8610–8618.

21 M. L. Heuzé, O. Collin, E. Terriac and M. Piel, Cell Migration, Humana Press, Totowa, NJ, 2011, vol. 769.

22 R. N. Palchesko, L. Zhang, Y. Sun and A. W. Feinberg, PLoS One, 2012, 7, e51499.

23 T. G. Kuznetsova, M. N. Starodubtseva, N. I. Yegorenkov, S. A. Chizhik and R. I. Zhdanov, Micron, 2007, 38, 824–833.

24 T. Iskratsch, H. Wolfenson and M. P. Sheetz, Nat. Publ. Gr., 2014, 15, 825–833.

25 E. Evans and B. Kukan, Blood, 1984, 64, 1028–1035.

26 O. Campàs, T. Mammoto, S. Hasso, R. a Sperling, D. O’Connell, A. G. Bischof, R. Maas, D. a Weitz, L. Mahadevan and D. E. Ingber, Nat. Methods, 2013, 11, 183–189.

27 L. Trichet, O. Campàs, C. Sykes and J. Plastino, Biophys. J., 2007, 92, 1081–9.

28 T. Mason and J. Bibette, Phys. Rev. Lett., 1996, 77, 3481–3484.

29 A. Millius and O. D. Weiner, Methods Mol. Biol., 2010, 591, 147–58.

30 P. Vargas, E. Terriac, A.-M. Lennon-Duménil and M. Piel, J. Vis. Exp., 2014, e51099.

31 J. Schindelin, I. Arganda-Carreras, E. Frise, V. Kaynig, M. Longair, T. Pietzsch, S. Preibisch, C. Rueden, S. Saalfeld, B. Schmid, J.-Y. Tinevez, D. J. White, V. Hartenstein, K. Eliceiri, P. Tomancak and A. Cardona, Nat. Methods, 2012, 9, 676–82.

32 A. Yeung, T. Dabros, J. Masliyah and J. Czarnecki, Colloids Surfaces A Physicochem. Eng. Asp., 2000, 174, 169–181.

33 F. Leal-Calderon, V. Schmitt and J. Bibette, Emulsion Science: Basic Principles, Springer, 2nd ed., 2007.

34 K. Ben M’Barek, D. Molino, S. Quignard, M. Plamont, Y. Chen, P. Chavrier and J. Fattaccioli, Biomaterials, 2015, 51, 270–277.

35 J. Fattaccioli, J. Baudry, N. Henry, F. Brochard-Wyart and J. Bibette, Soft Matter, 2008, 4, 2434–2440.

36 K. A. Brakke, Exp. Math., 1992, 1, 141–165.

37 S. J. Collins, F. W. Ruscetti, R. E. Gallagher and R. C. Gallo, J. Exp. Med., 1979, 149, 969–74.

38 A. B. Hauert, S. Martinelli, C. Marone and V. Niggli, Int. J. Biochem. Cell Biol., 2002, 34, 838–854.

39 E. S. Wittchen, R. a. Worthylake, P. Kelly, P. J. Casey, L. a. Quilliam and K. Burridge, J. Biol. Chem., 2005, 280, 11675–11682.

40 K. Wilson, A. Lewalle, M. Fritzsche, R. Thorogate, T. Duke and G. Charras, Nat. Commun., 2013, 4, 2896.

41 P.-G. de Gennes, F. Brochard-Wyart and D. Quéré, Capillarity and Wetting Phenomena, Springer New York, New York, NY, 2004.

42 V. Niggli, FEBS Lett., 1999, 445, 69–72.

43 T. Ishizaki, M. Uehata, I. Tamechika, J. Keel, K. Nonomura, M. Maekawa and S. Narumiya, Mol. Pharmacol., 2000, 57, 976–983.

44 S. Tamilvanan, Prog. Lipid Res., 2004, 43, 489–533.

45 P. Than, L. Preziosi, D.. Josephl and M. Arney, J. Colloid Interface Sci., 1988, 124, 552–559.

46 W. H. Guilford, R. C. Lantz and R. W. Gore, Am. J. Physiol. Physiol., 1995, 37, C1308–C1312.

47 C. M. Kraning-Rush, J. P. Califano and C. a. Reinhart-King, PLoS One, 2012, 7.

48 V. D. Varner and C. M. Nelson, Integr. Biol. (Camb)., 2013, 5, 1162–73.

49 M. Prass, K. Jacobson, A. Mogilner and M. Radmacher, J. Cell Biol., 2006, 174, 767–72.

50 T. Harada, J. Swift, J. Irianto, J.-W. Shin, K. R. Spinler, A. Athirasala, R. Diegmiller, P. C. D. P. Dingal, I. L. Ivanovska and D. E. Discher, J. Cell Biol., 2014, 204, 669–682.

51 K. Wolf, M. Te Lindert, M. Krause, S. Alexander, J. Te Riet, A. L. Willis, R. M. Hoffman, C. G. Figdor, S. J. Weiss and P. Friedl, J. Cell Biol., 2013, 201, 1069–84.

